# Functional surrogacy enables Vascular Ehlers-Danlos Syndrome modelling in zebrafish in the absence of a *COL3A1* ortholog

**DOI:** 10.64898/2026.07.03.736265

**Authors:** Daniel A. Baird, Nataliya Pidlisnyuk, Anna Matischen, Zuzanna Matelowska, Sookyung Seo, Nurhaziqah Supari, Jessica Bowen, Glenda Sobey, Meena Balasubramanian

**Author notes:** Corresponding author: Meena Balasubramanian, The Bateson Centre for Disease Mechanisms, School of Medicine and Population Health, Faculty of Health, University of Sheffield, Sheffield, United Kingdom.

## Abstract

Pathogenic variants in *COL3A1* cause Vascular Ehlers-Danlos syndrome (vEDS), a rare connective tissue disorder characterised by vascular fragility, increasing the risk of arterial ruptures/dissection. Advances in genomic sequencing have led to an increasing number of *COL3A1* variants where the clinical significance is unclear, with these being termed ‘variants of uncertain significance’ (VUS). VUS creates challenges for diagnosis and clinical management. Thus major efforts have been made to reclassify these to either pathogenic or benign variants in disease causality. Functional data from model systems can provide significant evidence to clinicians on the pathogenicity of a variant. To address the increasing numbers of VUS in *COL3A1*, we developed a fast pipeline using F0 crispant zebrafish to provide functional evidence for variant classification despite there being no direct orthologue of *COL3A1* in zebrafish. Loss of *col5a1* resulted in cardiac defects, dysmorphic blood vessel structures and delayed angiogenic sprouting. Trunk haemorrhage prevalence under physical stress increased in *col5a1* knockout zebrafish, recapitulating vEDS patients. Remarkably, co-injection of F0 *col5a1* knockout crispants with human wildtype *COL3A1* mRNA partially rescued cardiac and vascular phenotypes, indicating a level of functional conservation between zebrafish type V and human type III collagen. These findings establish a tractable *in vivo* platform for functional assessment of *COL3A1* VUS. Phenotypic rescue with wildtype *COL3A1* provides a benchmark against which the pathogenicity of variants can be evaluated, generating functional evidence for VUS reclassification. Our model provides both a valuable tool for investigating vEDS disease mechanisms and a clinically relevant platform to improve diagnoses for patients with suspected vEDS.

## INTRODUCTION

Vascular Ehlers-Danlos syndrome (vEDS, OMIM #130050) is a rare connective tissue disorder caused by autosomal dominant pathogenic variants in *COL3A1* with an estimated prevalence of 1 in 50,000-200,000 individuals^1^. Patients with vEDS are primarily characterised by blood vessel fragility, resulting in a greater risk of arterial rupture/dissection, intestinal/uterine fragility and blood vessel rupture in hollow organs. Due to the severity of these ruptures and vascular events, the mean life expectancy for these patients is lower compared to that of the general population. vEDS patients can also present with easy bruising due to these fragile vessels^2, 3^.

The *COL3A1* gene encodes for type III collagen, a homo-trimeric protein consisting of three α1 (III) α-chain monomers and a repeating Glycine-Xaa-Yaa triple helix domain. The COL3A1 protein is found in high abundance within the extracellular matrix (ECM), particularly in the ECM of blood vessels, the dermis, and hollow organs. Protein abundance in these tissues highlights the critical role it plays in supporting structural integrity and elasticity of soft tissues. Defects to the structure and/or function of type III collagen can result in ECM instability, resulting in the vascular events observed in vEDS patients^4, 5^. The underlying disease mechanisms are not fully understood. However, it is believed that either defective secreted levels or accumulation of secreted malformed type III collagen protein could lead to ECM weakening in blood vessels, making them viable to rupture^4^.

Despite knowing *COL3A1* pathogenic variants result in vEDS, the diagnosis of this is challenging. vEDS diagnosis can be missed due to overlapping clinical phenotypes in other connective tissue disorders, such as Marfan and Loeys-Dietz syndromes^6^. Initial clinical diagnosis is often made based on family history of the disease and clinical phenotyping. However, about 50% of vEDS probands have no family history of vEDS. An absence of family history can result in a diagnosis only being made after a major vascular event. The clinical presentation of vEDS can also be heterogeneous, even in those with the same variant, making diagnosis difficult even for experienced clinicians^7, 8^.

Due to significant advances in next generation sequencing technologies, the number of variants being identified has dramatically increased in recent years. To determine whether these variants are causative of disease or not, a large amount of evidence is required to classify the pathogenicity of these variants, such as population frequency data, computational predictions, familial segregation studies, and functional data. In many cases evidence for pathogenicity is lacking, especially in rare disorders, resulting in these variants being classified as ‘variants of uncertain significance’ (VUS)^9^. A VUS diagnosis can complicate clinical decision-making, potentially leading to delayed surveillance and access to disease-modifying treatments but also unnecessary costs on the healthcare service^10^. Consequently, diagnostic laboratories have launched significant efforts to reclassify VUS to provide a true diagnosis for these patients. One way to reclassify a VUS is to develop functional evidence from either an *in vitro* or *in vivo* model system.

Animal models for VUS interpretation have increased in usage in recent years, with zebrafish (*Danio rerio*) becoming a popular model for this. Zebrafish are used extensively in biomedical research due to their optical transparency for live-imaging of organs, large clutch sizes allowing for high-throughput studies, high genetic similarity to humans, shared developmental processes and molecular mechanisms, and external fertilisation allowing for easy genetic manipulation^11^. Zebrafish have also recently been utilised effectively to provide functional evidence for VUS reclassification and clinical advice in skeletal disorders^12^, spinal muscular atrophy^13^, and cerebral small vessel disease^14, 15^. Animal models investigating null and missense variants in vEDS have improved our understanding of the disease^16–18^. These have primarily been achieved in mice where technical challenges, slow reproductive timescales, small sample sizes, and high costs hampers VUS evaluation at a high through-put scale. Patient-derived fibroblasts have also been used to extend our knowledge on vEDS pathology and disease mechanisms^19, 20^. However, these 2D *in vitro* models lack the three dimensional physiological environment with specific cell-cell interactions, meaning certain elements of vascular development cannot be modelled accurately^21^.

Zebrafish do not have a gene that encodes for type III collagen, which is an obvious limitation of using zebrafish for expanding our knowledge of vEDS pathology and disease mechanisms. Despite this, recent research has suggested that zebrafish type V collagen can be used as a surrogate model for investigating type III collagen. Zebrafish *col5a1* was found to be expressed in the ECM of blood vessel associated perivascular fibroblast cells, and mutations to *col5a1* resulted in similar clinical signs to those with vEDS^22^.

In this study, we generated a fast and efficient F0 crispant knockout (KO) model of *col5a1* to highlight the critical role type V collagen plays in cardiovascular development. Using this F0 KO model, we were able to co-inject human wildtype *COL3A1* (*COL3A1*^WT^) mRNA to encode for human type III collagen when *col5a1* was removed from the zebrafish genome. In doing so, we found that human *COL3A1*^WT^ mRNA did partially rescue vascular abnormalities in knockout zebrafish crispants for *col5a1*. Together, our work provides important insights into the functional redundancy between human *COL3A1* and zebrafish *col5a1*, and suggests that zebrafish can be used for providing functional evidence in VUS reclassification.

## METHODS

### Evolutionary gene conservation tree, amino acid similarity & scRNA sequencing analysis

Evolutionary conservation tree of the *COL3A1* gene showing the phylogenetic history of gene gain and loss was generated using the ‘Gene gain/loss tree’ section under ‘Comparative Genomics’ for human *COL3A1* (ENSG00000168542). A simplified version of this gene gain/loss tree was generated using the phyloT (https://phylot.biobyte.de/) and iTOL: Interactive Tree of Life softwares^23^. Amino acid similarity between COL1A2, COL3A1, and COL5A1 between human, mouse, rat, rabbit, guinea pig, chimpanzee, macaque, pig, chicken, and zebrafish datasets were calculated using Protein Blast^24^. Protein accession or UniProt codes used can be found in Supplementary Table 1. Re-analysis of single-cell RNA sequencing from zebrafish was conducted using datasets from the DanioCell Desktop Companion App and online resource^25^. Expression of *col1a2* and *col5a1* was calculated from cell clusters for vascular (hema.2, hema.7, hema.18, hema.24, hema.30, hema.31), sclerotome (musc.21), osteoblast (mese.31), and mesenchymal cartilage (mese.5, mese.7, mese.15) cells limited to the 2 day old stage. Graphs were plotted on the DanioCell Desktop Companion App.

### AlphaFold predictions

AlphaFold protein structure predictions were generated using the AlphaFold Server by inserting the relevant amino acid sequences for wildtype human, wildtype zebrafish, and mutant zebrafish^26^.

### Zebrafish husbandry & maintenance

Wildtype and *Tg(myl7*:*citrine,kdrl:mCherry)* transgenic zebrafish line stocks were held at the Sheffield Biological Services aquarium housed at the Bateson Centre for Disease Mechanisms at the University of Sheffield, UK. The aquarium used to house zebrafish is approved by the UK Home Office and maintained in accordance with the UK Animals (Scientific Procedures) Act 1986. Zebrafish were raised in a 14:10 hour light-dark cycle at 27-28°C according to standard protocols. Staging and maintenance of embryos and larvae were performed as previously described and according to standard protocols^27^. Experiments were all conducted according to the project licence PP5148348 approved by the UK Home Office.

### crRNA design & selection

crRNAs targeting zebrafish *col5a1* (ENSDARG00000012593) were designed according to previously described protocols^28^. Briefly, four crRNAs targeting exons within the *col5a1* gene were designed using online tools from IDT (Integrated DNA Technologies) with the aim of increasing the probability of introducing biallelic frameshift mutations. Selection of crRNAs were based on predicted off- and on-target score rankings using CCTOP (https://cctop.cos.uni-heidelberg.de:8043)^29^, CHOPCHOP (https://chopchop.cbu.uib.no/)^30^, and CRISPOR (http://crispor.tefor.net)^31^. Sequences of designed crRNAs, IDT design IDs, and targeted exons are shown in Supplementary Table 2.

### gRNA/Cas9 RNP complex assembly & microinjection

Assembly of the gRNA/Cas9 ribonucleoprotein (RNP) complexes for *col5a1* and *scramble* negative controls was performed as previously described^28^. Equimolar volumes of 200 µM crRNAs and 200 µM tracrRNA were diluted in nuclease-free Duplex buffer (IDT) and incubated for 5 minutes at 95°C. Equimolar volumes of crRNA:tracrRNA and Alt.R S.p. HiFi Cas9 Nuclease V3, 61 µM (IDT) were mixed and incubated at 37°C for 5 minutes. RNP complexes for each condition were then pooled together. For microinjection, approximately 0.5 nL of pooled RNP was microinjected into the one-cell stage of wildtype or *Tg(myl7:citrine,kdrl:mCherry)* embryos. Survival of injected embryos were monitored several hours after microinjection and at 1 day post fertilisation (dpf).

### Genotyping & Sanger sequencing

To check for genetic edits, genomic DNA from individual embryos either at 1 or 2 dpf were initially taken by extraction with 50 mM NaOH at 95°C for 30 minutes. Samples were then neutralised with 1:10 1M Tris-HCl (pH 8). Polymerase chain reaction (PCR) was performed and samples ran for gel electrophoresis on a 3% agarose gel. For Sanger sequencing at 2 dpf, PCR products were purified using ExoSAP-IT reagents followed by Sanger sequencing conducted by Azenta GENEWIZ. Sequencing results were then analysed using Synthego ICE (Inference of CRISPR Edits)^32^ and TIDE (Tracking of Indels by DEcomposition)^33^ online tools. Primer sequences used are listed in Supplementary Table 3. Once it was determined that > 90% of injected samples had genetic edits, samples were genotyped in batches of 10 post experimental protocols.

### Quantitative Real-Time PCR (qRT-PCR)

For each genotype, between 10-15 embryos at 2 dpf were pooled and total RNA extracted using Trizol and Direct-zol RNA MiniPrep kit (Zymo Research). Each experimental group had three biological replicates. Total RNA extracted was used to synthesise random-primed cDNA with Reliance Select cDNA Synthesis kit (BioRad). Resultant cDNA was amplified using iTaq Universal SYBR Green Supermix (BioRad) using the BioRad CFX96 Real-Time System. The normalised relative expression for these genes was calculated against *rpl13a* expression. Primers for qRT-PCR were designed to target the end of the sequence and are listed in Supplementary Table 3.

### Phenotyping of F0 crispants

Phenotypic deviations of crispants were determined at 2 and 5 dpf, with samples categorised into four groups based on external visible features. Healthy zebrafish without any physical alterations from the normal state were classified as ‘no defect’. Zebrafish with a small amount of fluid accumulation in the pericardium were classified as ‘mild cardiac edema’, with a greater amount of accumulated fluid impacting normal swimming ability classified as ‘severe cardiac edema’. Any other visible phenotypes that were changed from the norm were put into an ‘other defects’ category. Zebrafish were phenotyped across multiple different microinjecting experiments. Brightfield images were taken on a Zeiss AxioZoom v16.

### Heart rate quantification

Zebrafish at 2 and 5 dpf were acclimatised to room temperature in batches and 1:100 MS222 added to larvae at 5 dpf. Heart rates were measured manually for 15 seconds and beats per minute calculated. Measurements were then normalised to the average of uninjected samples per experiment.

### Blood vessel morphology assessment

Transgenic *Tg(kdrl:mCherry)* zebrafish at 2 and 5 dpf were imaged in the lateral orientation using a Zeiss AxioZoom v16 fluorescent microscope. To measure the length of the intersegmental blood vessels, a previously published ImageJ-based method was utilised to determine the length of all blood vessels visible within captured images. An average blood vessel length per fish was then calculated from methods described previously^34^. Width measurements were performed on six intersegmental blood vessels per sample in each group, with the six vessels within the middle of the trunk measured. Measurements were taken at the thickest part of each blood vessel. An average blood vessel width from the six measurements for each sample was then calculated. For determination of the dorsal aorta width, six points along the entire length of the dorsal aorta for each sample was measured, before an average width of each dorsal aorta was calculated for each sample. Measurements were then normalised to the average of uninjected samples per experiment.

### Early developmental stage physical stress test

Physical stress test from 2 to 3.5 dpf was performed following previously described protocols^28^. For each experimental condition, two treatment options were possible. Half of each experimental group were placed in normal E3 media used for raising zebrafish under standard protocols, and half was raised in 0.6% methylcellulose made up in E3. Assessment of haemorrhage prevalence within the trunk was then performed at 3, 6, 24 and 27 hours post treatment. Samples were then culled at the end of the physical stress test.

### mRNA synthesis for rescue experiments

A full-length coding sequence of human *COL3A1*^WT^ (Genbank accession number NM_000090.4) was inserted into the pcDNA3.1(+) plasmid by GenScript. The plasmid was linearised with NotI before mRNA was *in vitro* transcribed to synthesise capped *COL3A1*^WT^ mRNA with mMessage mMachine T7 transcription kit (Invitrogen). Microinjections to generate crispants were performed as previously but with 0.5 nL of 25 pg *COL3A1*^WT^ mRNA included in the microinjection cocktail.

### Statistical analysis

Statistical analysis was performed using GraphPad Prism v10.6.1. Data in graphs are plotted as mean ± SEM unless indicated otherwise in the figure legend. For all statistical analyses, the level of significance was set at 0.05. Data was assessed for normal and lognormal distributions using the Shapiro-Wilk test. Statistical significance was determined using either ordinary one-way ANOVA or two-way ANOVA with Tukey’s multiple comparisons test, lognormal ordinary one-way ANOVA with Tukey’s multiple comparisons test, or Kruskal-Wallis test with Dunn’s multiple comparisons test, as detailed in figure legends. *P-*values are represented as follows: not significant (ns), *\*p* < 0.033, \*\**p* < 0.002, \*\*\**p* < 0.001.

## RESULTS

### Zebrafish *col5a1* selected as a surrogate gene replacement for human *COL3A1*

As zebrafish lack a direct orthologue of human *COL3A1*, as do C. elegans and fruit flies (Fig. 1A), we evaluated candidate fibrillar collagen genes previously proposed as alternatives for vEDS modelling - *col1a2* and *col5a1*^22^. Comparative protein analyses showed that zebrafish Col5a1 shared a greater high level of amino acid conservation to vertebrate species utilised in biomedical and agricultural research, as well as humans, than zebrafish Col1a2 does (Fig. 1Bi & ii). Additional analysis revealed that Col5a1 exhibited broad similarity to type III collagen proteins from humans and multiple other vertebrate species, as does Col1a2 (Fig. 1Biii & iv). Alignment of the C-terminal propeptide domain of human COL3A1 and zebrafish Col5a1 revealed 41% amino acid identity and 55% chemical similarity.

**Figure 1.**
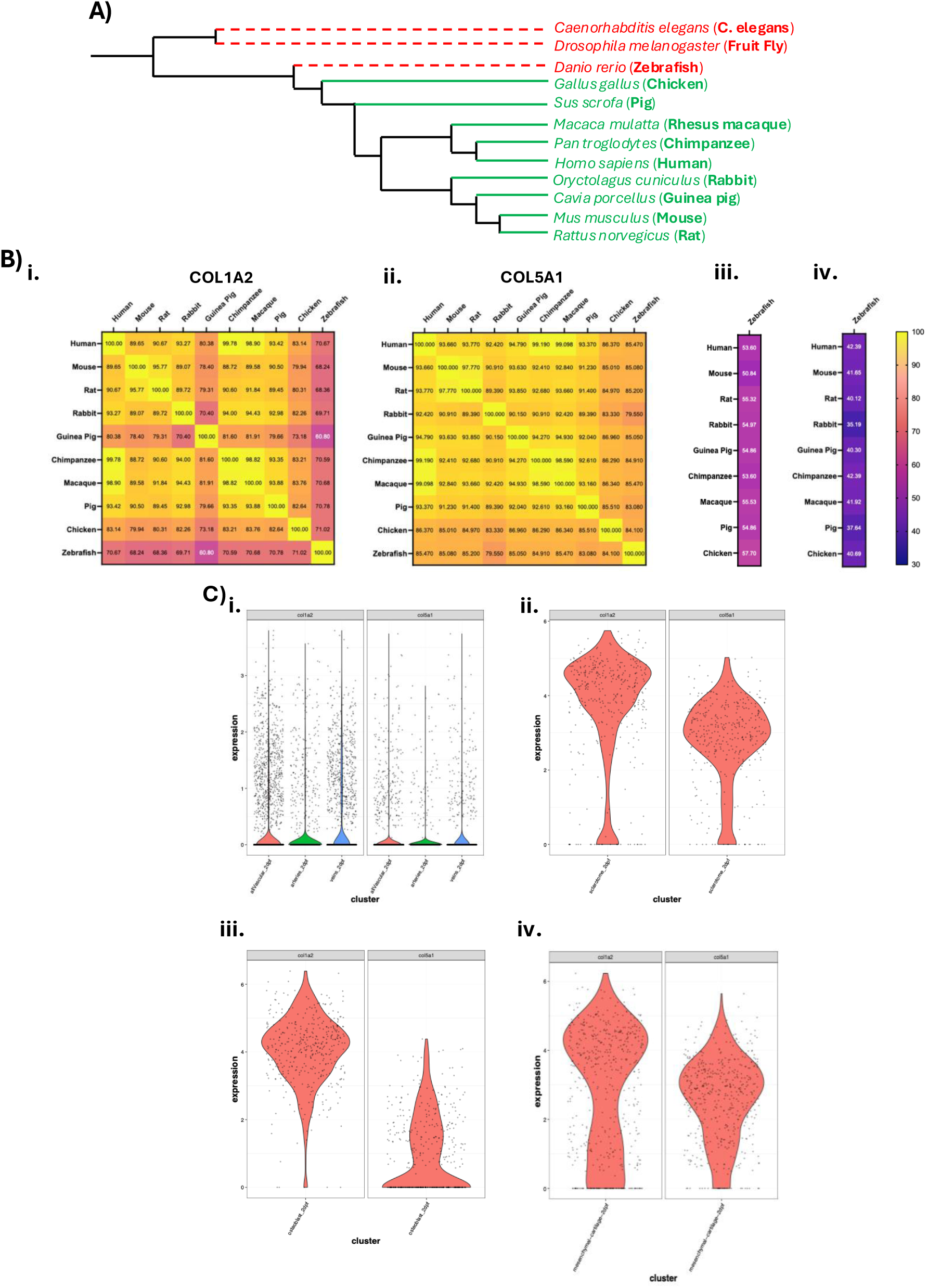
Protein conservation and RNA expression levels of fibrillar collagens. **A.** Evolutionary and protein conservation of type III collagen between human and commonly used model systems in biomedical and agricultural research. Phylogenetic tree showing gene conservation between species with species split over time. Green lines and text display conservation of type III collagen. Red dotted lines and text display gene loss and lack of type III collagen conservation. **B.** Protein amino acid similarities between humans and commonly used model systems between **i)** COL1A2, **ii)** COL5A1 and amino acid similarities between humans plus other commonly used model systems against zebrafish **iii)** Col1a2 and **iv)** Col5a1. Inner text shows percentage amino acid similarity between species. **C.** Violin plots showing expression (log2) at 2 dpf of *col1a2* (right side) and *col5a1* (left side) in **i)** vasculature clusters (red - all vascular cells, green - arterial cells, blue - venous cells), **ii)** sclerotome, **iii)** osteoblasts, and **iv)** mesenchymal cartilage clusters.

To further assess gene suitability, we examined publicly available single-cell RNA sequencing data from the DanioCell atlas. Both *col1a2* and *col5a1* were expressed similarly in vascular cell populations at 2 dpf, although expression of *col1a2* was higher in arterial cells (Fig. 1Ci). Expression of *col1a2* was more substantially enriched in the sclerotome, osteoblast, and mesenchymal cartilage cell populations at 2 dpf relative to *col5a1* (Fig. 1Cii-iv). Amongst the zebrafish type V collagen genes, *col5a1* was the consistently higher than the other transcripts in the sclerotome, osteoblasts, and mesenchymal cartilage, with a similar expression profile between *col5a1* and *col5a2a* in the vasculature at 2 dpf (Suppl. Fig. 1). Based on its protein conservation, vascular expression, and reduced enrichment in developing skeletal and cartilage cell populations, *col5a1* was selected as the surrogate gene for the generation of a zebrafish vEDS model.

### Targeting multiple exons simultaneously generates F0 crispant knockout for *col5a1*

To investigate the functional role of *col5a1* in vascular development, we utilised a fast and effective F0 knockout approach by targeting multiple exons of the *col5a1* gene simultaneously to introduce indels that cause a biallelic frameshift in the open reading frame. As *col5a1* has 61 exons, we targeted four distinct exons - exons 2, 3, 7 and 61 - with CRISPR-Cas9 guide RNAs. These exons encode for regions of the Col5a1 protein that are highly, but not completely, conserved between humans and mice (Fig. 2A). Sanger sequencing confirmed edits within the targeted exons, with representative sequencing chromatograms shown in Figure 2B. Gel electrophoresis of F0 crispant samples at 2 dpf shows multiple bands and smears, highlighting the presence of genetic edits (Suppl. Fig. 2).

**Figure 2.**
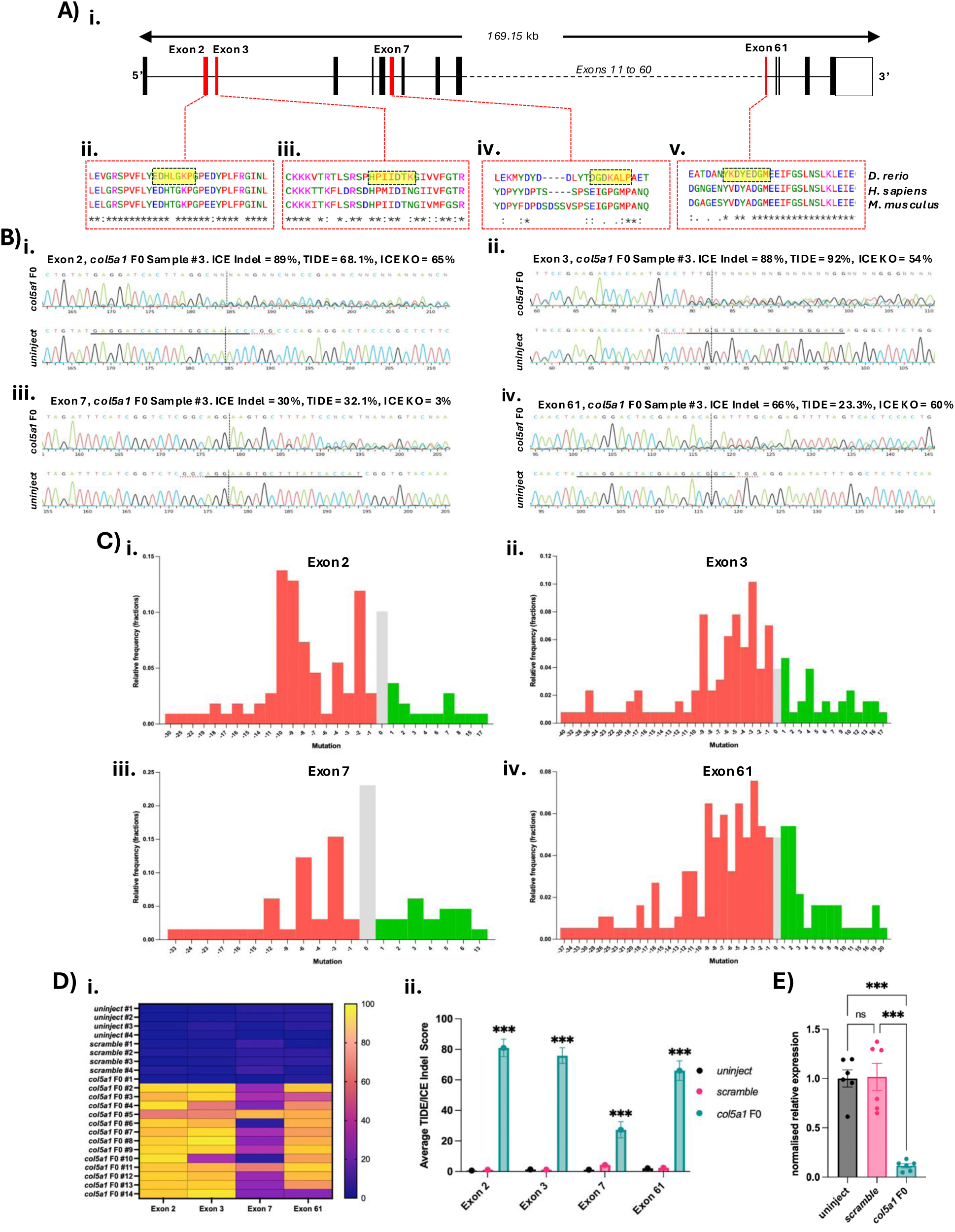
Generation of a *col5a1* F0 crispant knockout model. **A.** Schematic of *col5a1* F0 crispant targeting strategy. **i.** Schematic of *col5a1* gene with exon targets in red. Red boxes show amino acid sequences in targeted exonic regions with yellow highlights showing specific amino acids targeted by CRISPR-Cas9 reagents. Amino acid similarities between zebrafish (*D. rerio*), human (*H. sapiens*), and mice (*M. musculus*) are shown. ‘*’ represents fully conserved amino acid, ‘:’ represents highly conserved amino acid, and ‘.’ represents weakly conserved amino acid. Targets are shown for **ii)** exon 2, **iii)** exon 3, **iv)** exon 7, and **v)** exon 61. **B.** Representative Sanger sequencing chromatograms for **i)** exon 2, **ii)** exon 3, **iii)** exon 7, and **iv)** exon 61 with black solid line showing CRISPR-Cas9 target sequence, and dotted black line highlighting cut site in specific sequence. ICE indel, TIDE, and ICE KO scores are shown for specific examples. **C.** Relative frequency histogram for specific indels identified from Sanger sequencing in **i)** exon 2, **ii)** exon 3, **iii)** exon 7, and **iv)** exon 61. Red bars represent deletions. Green bars represent insertions. Grey bar represents no mutation. **D.** Indel scores for each exon target. **i)** Heat map showing average indel score for individual fish for individual exon. **ii)** Average indel score for each exon between uninjected, *scramble*, and *col5a1* F0 crispant embryos at 2 dpf. Two-way ANOVA with Tukey’s multiple comparisons test performed. **E.** Normalised relative expression of *col5a1* between experimental groups. One-way ANOVA with Tukey’s multiple comparisons test performed. ns, not significant. *** *p* < 0.001.

Using Synthego ICE and TIDE software to analyse Sanger sequencing results, we determined the relative frequency of specific insertions and deletions (indels) within each exon (Fig. 2C). We found that a 10 base pair (bp) deletion was the most frequent indel within exon 2 (13.8%) of injected *col5a1* F0 embryos. The most common indel in exons 3 and 61 was a 3 bp deletion, with a 10.2% and 7.7% frequency respectively. However, although indels were present within exon 7, the most frequent event was no indel present (23.1%). Analysis of protein structure by AlphaFold showed that wildtype zebrafish and human type V collagen proteins showed similar structures, despite AlphaFold having low confidence of prediction of these proteins (Suppl. Fig. 3A & B). However, when the most frequent indels in each exon are introduced and modelled, AlphaFold predicts a heavily truncated protein without a triple glycine helix domain (Suppl. Fig 3C)

Analysing individual zebrafish embryos at 2 dpf, it was clear that our CRISPR-Cas9 targeting reagents were highly effective in exons 2, 3, 61, with between 20-40% efficiency in exon 7 (Fig. 2Di). Although the editing efficiency in exon 7 was lower, the average indel score across all exons was significantly higher in those injected with gene editing reagents targeting *col5a1* than in controls (Fig. 2Dii).

To assess whether targeting multiple exons simultaneously does result in a gene knockout preventing protein transcription, we performed qRT-PCR targeting a region downstream of our exonic targets towards the 3’ end of the *col5a1* gene. We found that in *col5a1* F0 injected embryos, the normalised relative expression of *col5a1* was decreased to 0.12. This was significantly lower than uninjected and *scramble* controls, where the *col5a1* expression did not differ between the two controls (Fig. 2E). Zebrafish *col5a1* F0 crispants will now be referred to as *col5a1* knockouts (KOs).

### Defective cardiovascular function and development observed in *col5a1* KOs

To initially determine whether our *col5a1* KOs presented with any clear and obvious cardiovascular clinical signs, we performed phenotyping experiments at 2 dpf and found that *col5a1* KOs do present with varying levels of pericardial oedema, with limited amounts of phenotypic defects seen within uninjected and *scramble* control samples (Fig. 3A & Ci). Overall, a mean of 70.55% of *col5a1* KO embryos presented with a cardiac defect, compared to only 6.6% and 11.9% of uninjected and *scramble* injected embryos respectively. Furthermore, 93.4% of uninjected embryos and 87.75% of *scramble* injected embryos were found to have no cardiac defect, whereas only 29.45% of *col5a1* KOs had no cardiac defect (Fig. 3Cii). The prevalence of cardiac defects found in *col5a1* KOs was significantly greater compared to controls.

**Figure 3.**
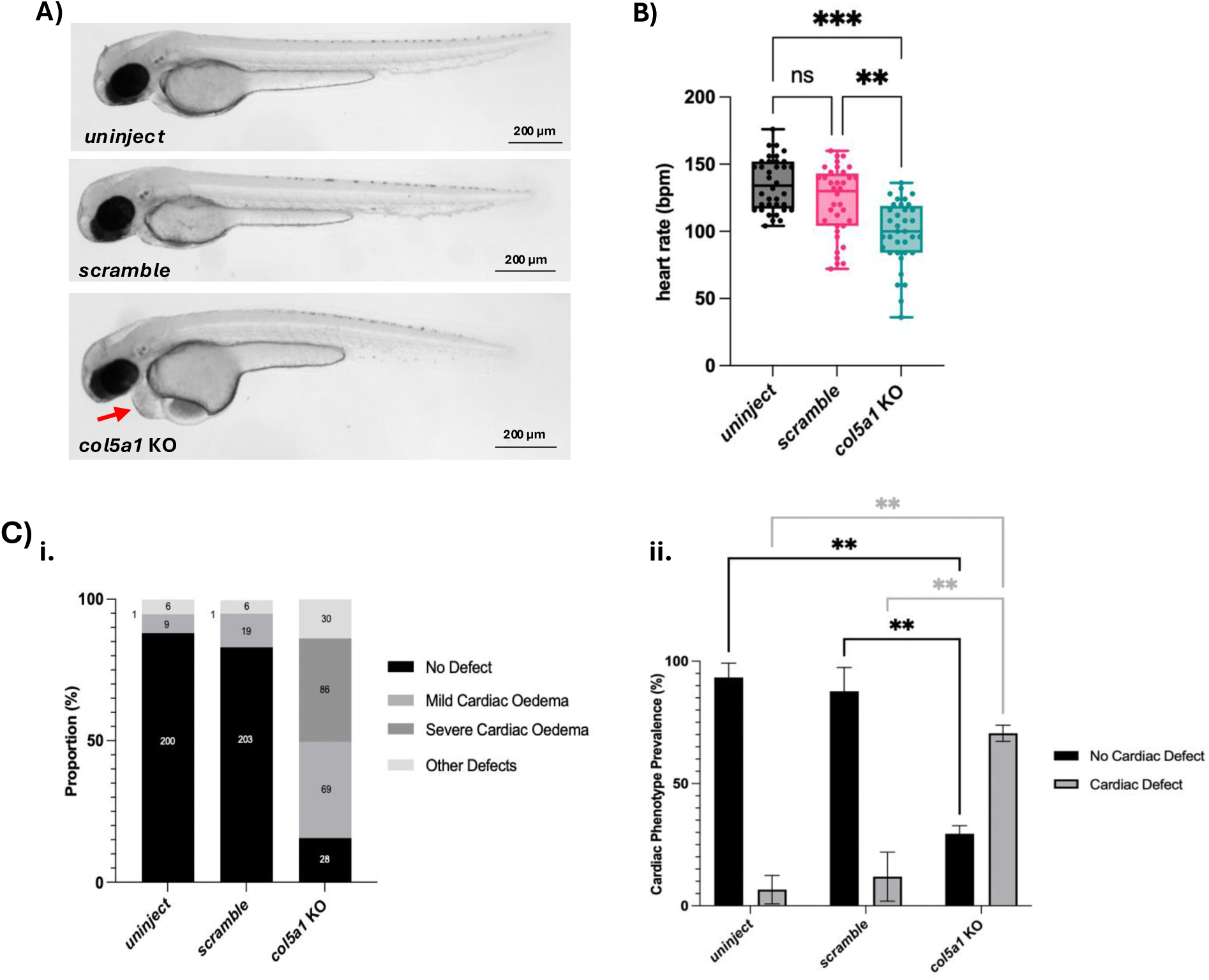
Cardiovascular phenotypes of *col5a1* F0 KOs at 2 dpf. **A.** Gross morphology brightfield images of 2 dpf uninject, *scramble*, and *col5a1* KO embryos with pericardial oedema highlighted by red arrow. Scale bars, 0.2 mm. **B.** Heart rate quantification of experimental embryos at 2 dpf. Kruskal-Wallis test with Dunn’s multiple comparisons test performed. **C.** Quantification of cardiac phenotypes observed in 2 dpf embryos. **i)** Proportion of embryos with differing levels of cardiac oedemas and other phenotypes. Embryo numbers shown in the middle of bars. **ii)** Prevalence of overall cardiac phenotypes observed between experimental groups. Two-way ANOVA with Tukey’s multiple comparisons test performed. ** *p* < 0.002. *** *p* < 0.001.

As the presence of pericardial oedema is indicative of a functional cardiac defect, we analysed the heart rate of 2 dpf embryos to assess whether knocking out *col5a1* disrupts cardiovascular function. It was found that the mean normalised heart rate for *col5a1* KOs was significantly slower when compared to both controls (Fig. 3B). We also observed that the normalised heart rate was significantly lower for *col5a1* KOs at 5 dpf (Suppl. Fig. 4A). No differences in normalised heart rates at either 2 or 5 dpf was found between uninjected and *scramble* injected controls.

### Blood vessel morphology and development of col5a1 KOs is disrupted

To assess the impact of removing the *col5a1* gene on vascular development, we analysed multiple morphological features of developing blood vessels using F0 KOs in the *Tg(kdrl:mCherry)* background (Fig. 4A). Although phenotypic differences in blood vessel structure were not obviously visualised at 2 dpf (Fig. 4B) or 5 dpf (Suppl. Fig. 4B), morphological measurements found that the length of intersegmental vessels (ISVs) in *col5a1* KOs were significantly smaller than that of uninjected and *scramble* controls at 2 dpf. Additionally, the width of the ISVs and the dorsal aorta at 2 dpf in *col5a1* KOs were both significantly smaller when compared to both controls (Fig. 4C). Smaller ISV lengths and ISV and dorsal aorta widths were also observed to be significantly at 5 dpf in *col5a1* KOs compared to control samples (Supp. Fig. 4C).

**Figure 4.**
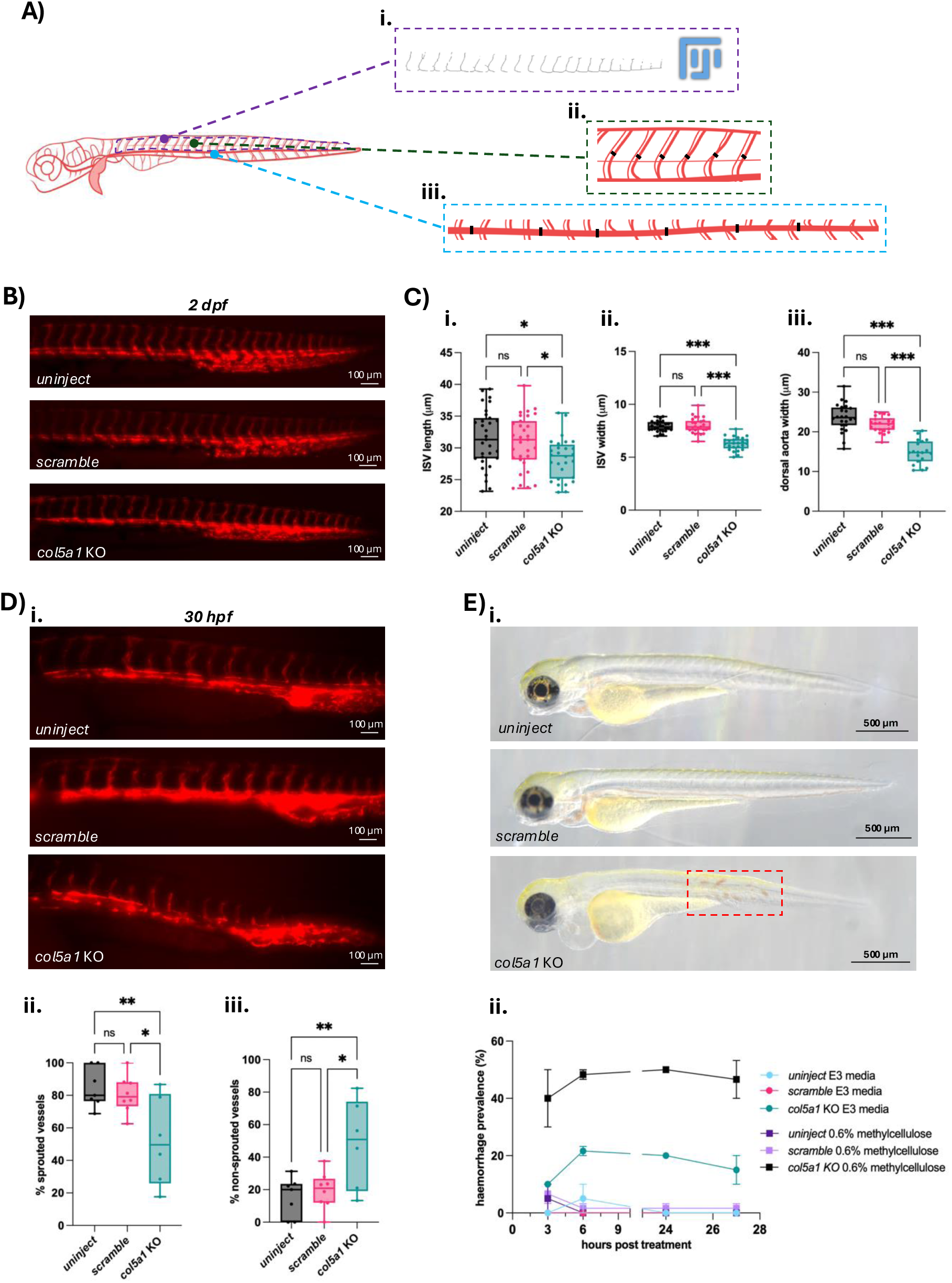
*col5a1* KO zebrafish display dysmorphic vessel phenotypes and vascular fragility. **A.** Schematic showing how blood vessel morphology was measured. **i)** Intersegmental blood vessel length was analysed using an ImageJ macro. **ii)** An intersegmental blood vessel width per sample was obtained by taking six measurements from the thickest point of vessels around the middle somite of the fish and an average of these taken to get an average width per sample. **iii)** Dorsal aorta widths were measured by taking six width measurements along the length of the vessel and averaged per sample. Created with BioRender **B.** Representative immunofluorescent images of *Tg(kdrl:mCherry)* positive samples for uninject, *scramble*, and *col5a1* KO embryos at 2 dpf. Scale bars, 0.1 mm. **C.** Quantification of average **i)** ISV length, **ii)** ISV width, and **iii)** dorsal aorta width in each experimental condition at 2 dpf. Each dot in **(i)** represents the average of the total number of vessels found in the embryo tested. Each dot in **(ii)** and **(iii)** represents an average of 6 vessel measurements in an embryo. Ordinary one-way ANOVA with Tukey’s multiple comparisons test performed. ns, not significant. * *p* < 0.033. *** *p* < 0.001. **D. i)** Representative immunofluorescent images of *Tg(kdrl:mCherry)* positive samples for uninject, *scramble*, and *col5a1* KO embryos at 30 hpf. Scale bars, 0.1 mm. Quantification of proportion of **ii)** sprouted and **iii)** non sprouted ISVs at 30 hpf. Ordinary one-way ANOVA with Tukey’s multiple comparisons test performed. ns, not significant. * *p* < 0.033. ** *p* < 0.002. **E. i)** Representative brightfield images of larvae 27 hours post 0.6% methylcellulose treatment (3.5 dpf) for early developmental physical stress tests. Red box highlights trunk haemorrhages in *col5a1* KO larvae. Scale bars, 0.5 mm. **ii)** Quantification of trunk haemorrhage prevalence in uninject, *scramble*, and *col5a1* KO zebrafish under either no physical stress or physical stress.

Blood vessels sprouting from the dorsal aorta, or angiogenesis, is a crucial part of ISV development. To determine whether this process is affected by knocking out *col5a1*, we assessed whether there were clear differences in ISV angiogenesis at 30 hours post fertilisation (hpf). A large proportion of ISVs in uninjected (84.56%) and *scramble* injected (80.64%) were fully sprouted at this timepoint, which was not observed in *col5a1* KO samples (51.86%). The proportion of ISVs not fully sprouted was significantly greater in *col5a1* KOs (48.31%) compared to uninjected and *scramble* controls (15.44% and 19.36% respectively) (Fig. 4D).

### Physical stress leads to increase in haemorrhage prevalence in col5a1 KOs

Since vEDS patients are characterised by blood vessel fragility, we investigated whether our *col5a1* KOs also presented with fragile blood vessels. This was done by raising samples at 2 dpf for 27 hours in 0.6% methylcellulose where the fish need to exert more energy to swim. Increasing the physical strength needed to swim resulted in haemorrhages observed in the trunk of *col5a1* KO fish 25 hours post treatment (hpt) (Fig. 4Ei). Trunk haemorrhage in *col5a1* KOs under physical stress was observed far more frequently when compared to uninjected and *scramble* control fish under physical stress. Furthermore, more trunk haemorrhages were observed in *col5a1* raised under physical stress than those not under any physical stressors. Little differences in prevalence were seen between control samples raised under physical stress compared to those not (Fig. 4Eii).

### Co-injection of wildtype COL3A1 mRNA partially rescues col5a1 KO phenotypes

To investigate whether zebrafish *col5a1* can be utilised as an effective surrogate gene for human *COL3A1*, we generated mRNA encoding for wildtype human *COL3A1* (*COL3A1*^WT^) and co-injected this into *col5a1* KO crispants. Phenotypic analysis at 2 dpf showed that those co-injected with *COL3A1*^WT^ mRNA did not present with cardiac oedemas as observed in *col5a1* KOs alone (Fig. 5A). Quantifying this, 35.39% of *col5a1* KOs presented with no cardiac defects compared to 72.60% of KOs rescued with *COL3A1*^WT^ with no cardiac abnormalities. 87.8% and 88.39% of uninjected and *scramble* samples at 2 dpf had no cardiac phenotype respectively. Furthermore, the proportion of embryos with a cardiac defect or other phenotypic abnormality was greater in *col5a1* KOs compared to *COL3A1*^WT^ co-injected, uninjected and *scramble* injected embryos (Fig. 5Bi).

**Figure 5.**
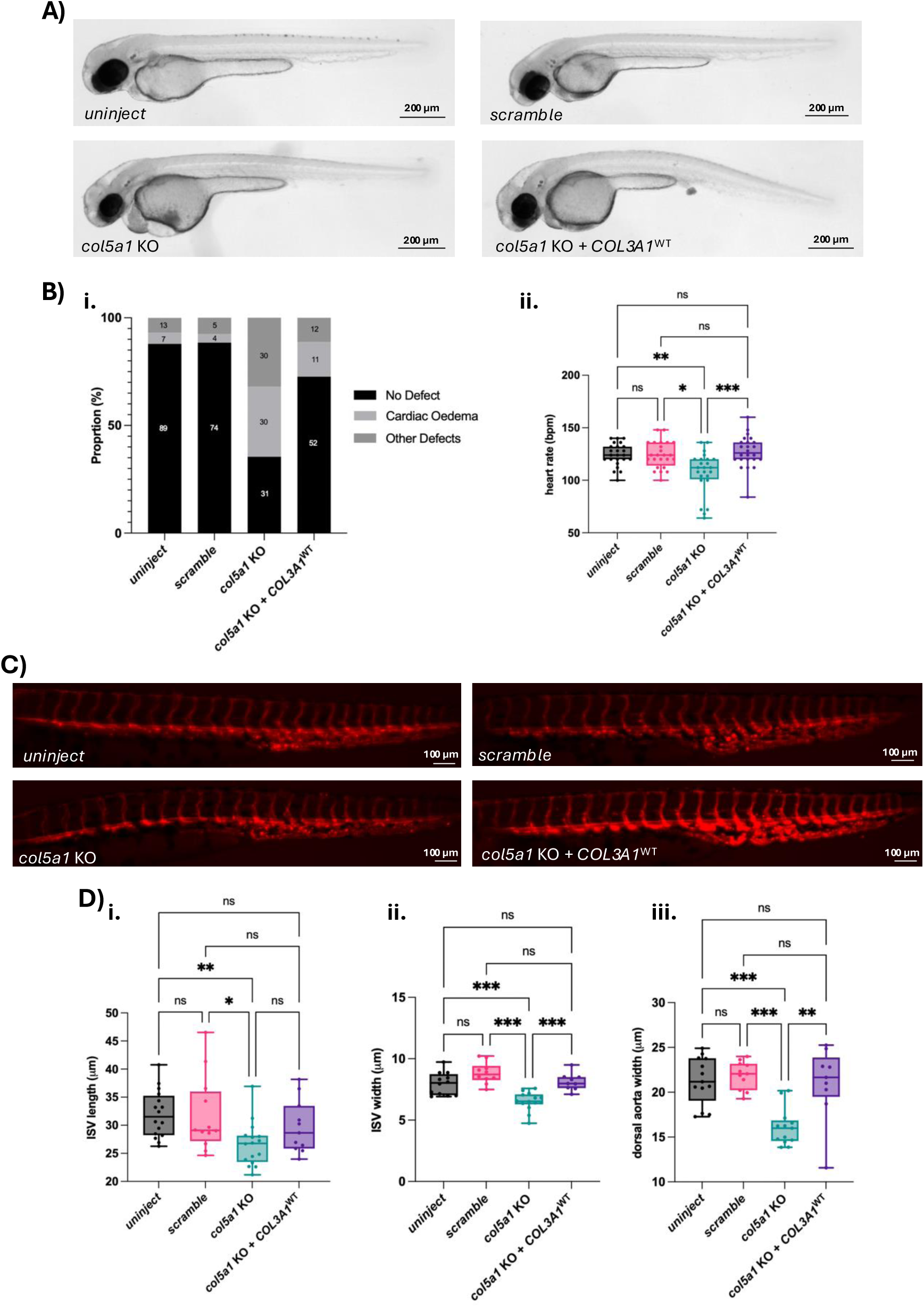
Partial functional rescue of cardiovascular phenotypes by human *COL3A1*^WT^ mRNA co-injection into *col5a1* KO crispants. **A.** Gross morphology brightfield images at 2 dpf of uninject, *scramble*, *col5a1* KO and KO co-injected with *COL3A1*^WT^ mRNA. Scale bars, 0.2 mm. **B.** Cardiovascular phenotype quantification. **i)** Proportion of cardiac oedema phenotypes in each experimental condition with embryo numbers shown in the middle of bars. **ii)** Heart rate quantification measurements at 2 dpf. Kruskal-Wallis test with Dunn’s multiple comparison test performed. ns, not significant. *\* p* < 0.033. ** *p* < 0.002. *** *p* < 0.001. **C.** Representative immunofluorescent images of *Tg(kdrl:mCherry)* positive samples for uninject, *scramble*, *col5a1* KO and KO co-injected with *COL3A1*^WT^ embryos at 2 dpf. Scale bars, 0.1 mm. **D.** Quantification of average **i)** ISV length, **ii)** ISV width, and **iii)** dorsal aorta width in each experimental condition at 2 dpf. Each dot in **(i)** represents the average of the total number of vessels found in the embryo tested. Each dot in **(ii)** and **(iii)** represents an average of 6 vessel measurements in an embryo. Lognormal ordinary one-way ANOVA with Tukey’s multiple comparisons test performed for **(i)**. Ordinary one-way ANOVA with Tukey’s multiple comparisons test performed for **(ii)**. Kruksal-Wallis test with Dunn’s multiple comparisons test performed for **(iii**). ns, not significant. *\* p* < 0.033. ** *p* < 0.002. *** *p* < 0.001.

When evaluating the heart rate at 2 dpf, it was found that the normalised heart rate of co-injected embryos with *COL3A1*^WT^ mRNA had significantly faster normalised heart rates when compared to *col5a1* KOs, but no difference in the rate was observed between mRNA co-injected fish and controls. KOs for *col5a1* still presented with slower normalised heart rates than control embryos (Fig. 5Bii).

Visualisation of transgenic ISVs between controls, KOs, and KOs with *COL3A1*^WT^ mRNA co-injection was performed (Fig. 5C). We found that at 2 dpf, the majority of those co-injected with *COL3A1*^WT^ mRNA did not present with the vascular morphological abnormalities observed in *col5a1* KOs alone. Rescued embryos had ISV vessel lengths and widths comparable to that of uninjected and *scramble* embryos, as was the width of the dorsal aorta between these fish. Again, it was seen that the length and width measurements in *col5a1* KOs was significantly smaller than both controls (Fig. 5D).

## DISCUSSION

The number of VUS being identified in connective tissue disorders has been increasing over the last decade. Reclassification of a VUS to provide an accurate diagnosis to a patient requires substantial evidence from multiple sources, including functional evidence from a model system. Zebrafish have increasingly been used to generate functional evidence for variant reclassification^12–15^, as they have less physiological and technical challenges compared to *in vitro* models and are more cost effective than rodent models. A major limitation in using zebrafish to model vEDS has been the absence of a direct orthologue of human *COL3A1*. To overcome this, previous studies have suggested that alternative zebrafish fibrillar collagen genes, *col1a2* and *col5a1*, may have overlapping functions within the vasculature to human *COL3A1*^22^. In this study, we evaluated these candidate genes and identified *col5a1* as the most suitable surrogate gene for modelling *COL3A1* and vEDS in zebrafish to better understand this disorder and to improve genetic diagnoses in patients suspected to have this disorder.

### Selection of *col5a1* as a surrogate gene for vEDS disease modelling

Actinopterygii divergence during evolution is a plausible reason for the lack of type III collagen in zebrafish and other actinopterygian fish, which may have occurred alongside duplication of other fibrillar collagen genes. This genomic duplication may have resulted in other collagens substituting for type III collagen in these fish^35^. Both *col1a2* and *col5a1* are expressed within the vascular cell populations, consistent with having a role in blood vessel development and homeostasis. Our analysis of publicly available single-cell RNA sequencing data revealed that *col1a2* is more expressed within the sclerotome, osteoblasts, and mesenchymal cartilage than *col5a1* during early development. Zebrafish mutants for *col1a2* are widely reported to present with skeletal phenotypes which recapitulate collagenopathies in humans, but have normal cardiac structure and function at larval and adult stages^12, 36^. As such, targeting *col5a1* with a more restricted expression profile may produce phenotypes that are more specific to the vasculature and vEDS pathology than *col1a2*. Additional evidence for selecting *col5a1* came from comparative protein analyses, which suggest a greater overall similarity between zebrafish Col5a1 and fibrillar collagens from other model systems and humans than was seen for zebrafish Col1a2. Shared protein conservation can be indicative of shared function, but this is not always the case^37, 38^. Collectively, this all indicates that *col5a1* is the most appropriate surrogate gene for modelling vEDS vascular pathologies in zebrafish.

### Functional rescue by human *COL3A1* mRNA validates *col5a1* surrogacy

A central challenge in modelling vEDS in zebrafish is the absence of a direct orthologue of human *COL3A1*. Consequently, vEDS research has naturally been undertaken in organisms with this gene, with the majority of work being carried out in mice. Heterozygous knockout and a transgenic missense mouse model for *Col3a1* both presented with vascular fragility^16, 18^, with haploinsufficient mouse models of *Col5a1* also presenting with structural blood vessel anomalies without any rupture phenotypes^39^. Thus, both type III and V collagen both play a role in blood vessel development and homeostasis. As zebrafish *col1a2* mutants appear to lack a vascular phenotype, this adds to the argument of utilising *col5a1* as a surrogate gene given its known functions in the vascular system. Whilst previous studies have suggested that zebrafish *col5a1* may have overlapping functions within the vascular system^22^, it is still unclear whether a *col5a1* loss-of-function zebrafish could accurately and efficiently model vEDS caused by *COL3A1* pathogenic variants. Addressing this question was the primary objective of our study.

The key finding from our work is that co-injection of a *col5a1* knockout F0 model with human wildtype *COL3A1* mRNA was able to partially rescue the cardiovascular phenotypes observed in *col5a1* crispants. Restoration of vascular morphology following co-injection demonstrates that human *COL3A1* can, in part, compensate for a complete loss of type V collagen in zebrafish vasculature development. These findings reflect a surprising functional overlap between human type III and zebrafish type V collagen in the vasculature, and highlights that *col5a1* can act as a surrogate gene for *COL3A1*.

The partial rescue is of note due to the overall modest amino acid similarity between the two proteins. Although not overly similar in terms of the specific amino acid sequence, the C-terminal propeptide domain of human type III and zebrafish type V collagens do share over 50% chemical similarity. This C-terminal domain is required for proper trimerisation and biosynthesis of collagen^40^. Conservation of biochemical properties within this domain may be sufficient to conserve elements of protein function, providing a possible explanation for the ability of human COL3A1 to compensate partially for the complete loss of zebrafish Col5a1.

### Clinical utility of *col5a1* model for *COL3A1* VUS reclassification

Our findings have important clinical implications. Variant interpretation remains a major challenge in not just vEDS, but in other EDS types. Increasing numbers of *COL3A1* variants classified as VUS are being seen following advances in next-generation sequencing technologies. Our findings establish a framework for future functional studies of *COL3A1* using zebrafish to address the challenge of increasing VUS diagnoses. Human mRNA encoding for known pathogenic and benign variants can be co-injected into the *col5a1* crispants to assess for rescue efficacy. Our work also paves the way for *COL3A1* VUS mRNA co-injection into *col5a1* knockout crispants and assessing whether vascular phenotypes are rescued or not. This would provide clinicians with functional evidence for variant reclassification efforts.

Phenotypes quantified in our study provide multiple readouts that can be used in future variant analysis pipelines. The partial rescue achieved with wildtype *COL3A1* mRNA also provides a biological benchmark against which VUS rescues can be compared to. Similar variant analysis approaches using zebrafish have recently been performed for osteogenesis imperfecta^12^, spinal muscular atrophy^13^, and cerebral small vessel disease^14, 15^. To our knowledge, our work is the first to perform this evaluation for vEDS and using a surrogate gene for an non-conserved gene between species.

### Mechanisms underlying vascular abnormalities by *col5a1* loss of function

Our study contributes additional knowledge to the important role of *col5a1* in the cardiovascular development and pathology. Loss of *col5a1* resulted in an increase in cardiac defects, bradycardia, and dysmorphic blood vessel structures. Biomechanical forces are known to be critical to various aspects of cardiac^41, 42^ and vascular growth and formation^43^, with altered blood flow through differing heart rates influencing vessel remodelling. An increased heart rate can promote vessel dilation^44^, whereas a slower heart rate reducing blood flow can disrupt ISV remodelling^45^. Bradycardia observed in *col5a1* crispants may contribute to vascular defects by altering flow mechanical forces. However, as type V collagen regulates fibrillogenesis and ECM development, vascular abnormalities observed may occur independently of blood flow^46, 47^.

Evidence supporting blood flow independent mechanisms comes from a study investigating *plxnd1*. Zebrafish mutants for *plxnd1* required induction of Col5a1 expression in vascular endothelial cells for hyperplastic growth in visceral adipose tissue, with a Col5a1-dependent increase in fibrillogenesis observed^48^. In our *col5a1* KO model, loss of Col5a1 would be expected to reduce fibrillogenesis within the developing vasculature. This may result in reduced ISV lengths and widths seen in crispants, due to this constituent being missing from the ECM. As the ECM plays a critical role in determining heart geometry and chamber development^49^, defects in ECM assembly may also contribute to the slower heart rate observed in *col5a1* crispants. Furthermore, lumenisation of the dorsal aorta, as well as other critical vessels in zebrafish, are governed by blood flow independent mechanisms^50, 51^. Morphometric differences to the ISV and dorsal aorta are unlikely to be solely altered by haemodynamic forces. Defective ECM formation and altered mechanical forces are likely acting together, possibly in a negative feedback mechanism, to drive vascular abnormalities in our vEDS model.

ISVs are formed through a process of sprouting angiogenesis, whereby endothelial tip cells emerge from pre-existing vessels and migrate towards the roof of the zebrafish trunk, guided by interactions with the ECM^52^. Collagens are known to promote endothelial cell migration during this process^53, 54^. Type I collagen is thought to act as a scaffold to support cells undergoing angiogenesis^51^, with type IV collagen regulating blood vessel formation in the brain^55^. Given the role of type V collagen in regulating collagen fibril assembly, its loss may compromise ECM architecture and limit structural support for endothelial cell migration. Consistent with this hypothesis, selective pharmacological inhibition of cells producing collagen 1α2 disrupts collagen scaffolds and endothelial cell migration within the zebrafish caudal fin^56^. Type V collagen co-assembles with type I collagen to form fibrils during overall embryonic development and cardiovascular homeostasis^57, 58^. Subsequently, loss of type V collagen may not only impact its function but also the broader collagen network. A lack of interaction between types I and V collagen may alter ECM morphology and its mechanical properties, limiting its ability to act as a proper scaffold for angiogenesis. Such alterations affecting ECM composition provides a potential explanation for the delayed and defective angiogenesis observed in our *col5a1* crispant model.

Vascular fragility is the primary feature of vEDS, with disruption to the structure, function and development of vascular endothelial cells known to cause vascular fragility^59^. Increasing physical stress on *col5a1* knockout crispants lead to an increased incidence of trunk haemorrhages. Collagen fibrils in vEDS patients exhibit reduced tensile strength and are unstable, predisposing the vessels to rupture^60, 61^. Fragile blood vessels may arise due to changes in the interaction between types I and V collagens, with reduced type V collagen levels disrupting fibril structures^62^. Defective fibril structures may lead to vessels being weaker, leading to an increased likelihood of vessel rupture and tissue instability. Furthermore, analysis of vEDS patient fibroblast cells have altered viscoelasticity properties when compared to control cells^63^. This further supports a role for type V collagen in vascular ECM development and homeostasis, with disruption to the composition and biomechanical properties of the ECM in *col5a1* crispants reducing vessel integrity, increasing their susceptibility to haemorrhage.

### Broader applications for Ehlers-Danlos syndrome modelling

Although this study was designed to model vEDS, pathogenic variants in *COL5A1* are causative of Classical Ehlers-Danlos syndrome (cEDS)^64^. These patients are characterised by skin hyperextensibility and joint hypermobility^65^. As type V collagen regulates fibrillogenesis with type I collagen^66^, mutations which result in a reduction of type V collagen protein would lead to defective ECM composition, structure and function. Given the major role type V collagen seems to play in collagen development and function, our zebrafish model may have a broader utility beyond vEDS. While the vascular phenotypes observed support its use for studying vEDS, our model may also provide a valuable platform for investigating disease mechanisms and therapies for cEDS, as well as other collagen related disorders.

In conclusion, *col5a1* loss of function zebrafish model recapitulates vascular fragility clinical signs associated with vEDS. Crucially, these phenotypes can be partially rescued by human *COL3A1*, demonstrating functional conservation between the two different genes, and the generation of a new model for investigating *COL3A1* in health and disease. Thus, our zebrafish vEDS model can be used as a system for studying vEDS disease mechanisms in greater detail and for rapid functional evaluation of VUS for variant reclassification. Our findings suggest evolutionary compatibility between zebrafish *col5a1* and human *COL3A1*. This establishes a new utility for zebrafish in modelling genetic disorders where the human ortholog of interest is not directly conserved between species.

## ACKNOWLEDGEMENTS

We would like to thank Catherine Loynes for excellent assistance and expertise, Emily Noël and Corinna Snashall for use of the *Tg(myl7*:*citrine,kdrl:mCherry)* line, and all the Biological Services Aquaria Team for fish care and practical assistance. We are grateful and like to thank Annabelle’s Challenge for funding this project.

## AUTHOR CONTRIBUTIONS

MB conceived the study. MB and DB supervised experiments and analyses. DB and MB wrote the manuscript with input from all authors (NP, AM, ZM, SS, NS, JB, GS). DB, NP, AM, ZM, SS, and NS performed zebrafish experiments and analyses. NP, JB, GS and MB provided clinical input in relation to the project.

## SUPPLEMENTAL FIGURE LEGENDS

**Suppl. Figure 1.**
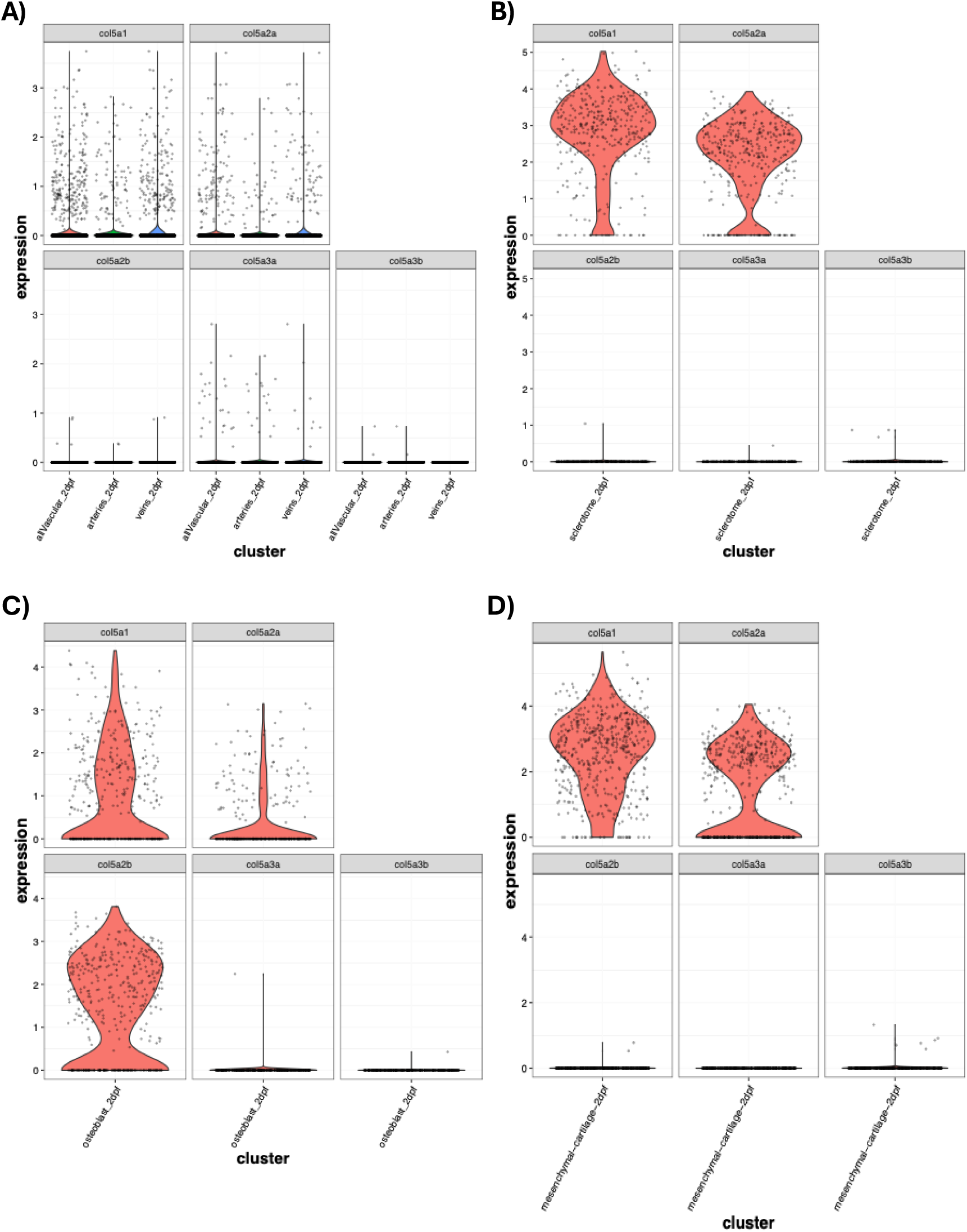
Expression levels of type V collagen zebrafish paralogs in different tissues. Violin plots showing expression (log2) at 2 dpf of *col5a1* (same as shown in Fig. 1), *col5a2a*, *col5a2b*, *col5a3a*, and *col5a3b* in **A.** vasculature clusters (red - all vascular cells, green - arterial cells, blue - venous cells), **B.** sclerotome cell clusters, **C.** osteoblast clusters, and **D.** mesenchymal cartilage cell clusters.

**Suppl. Figure 2.**
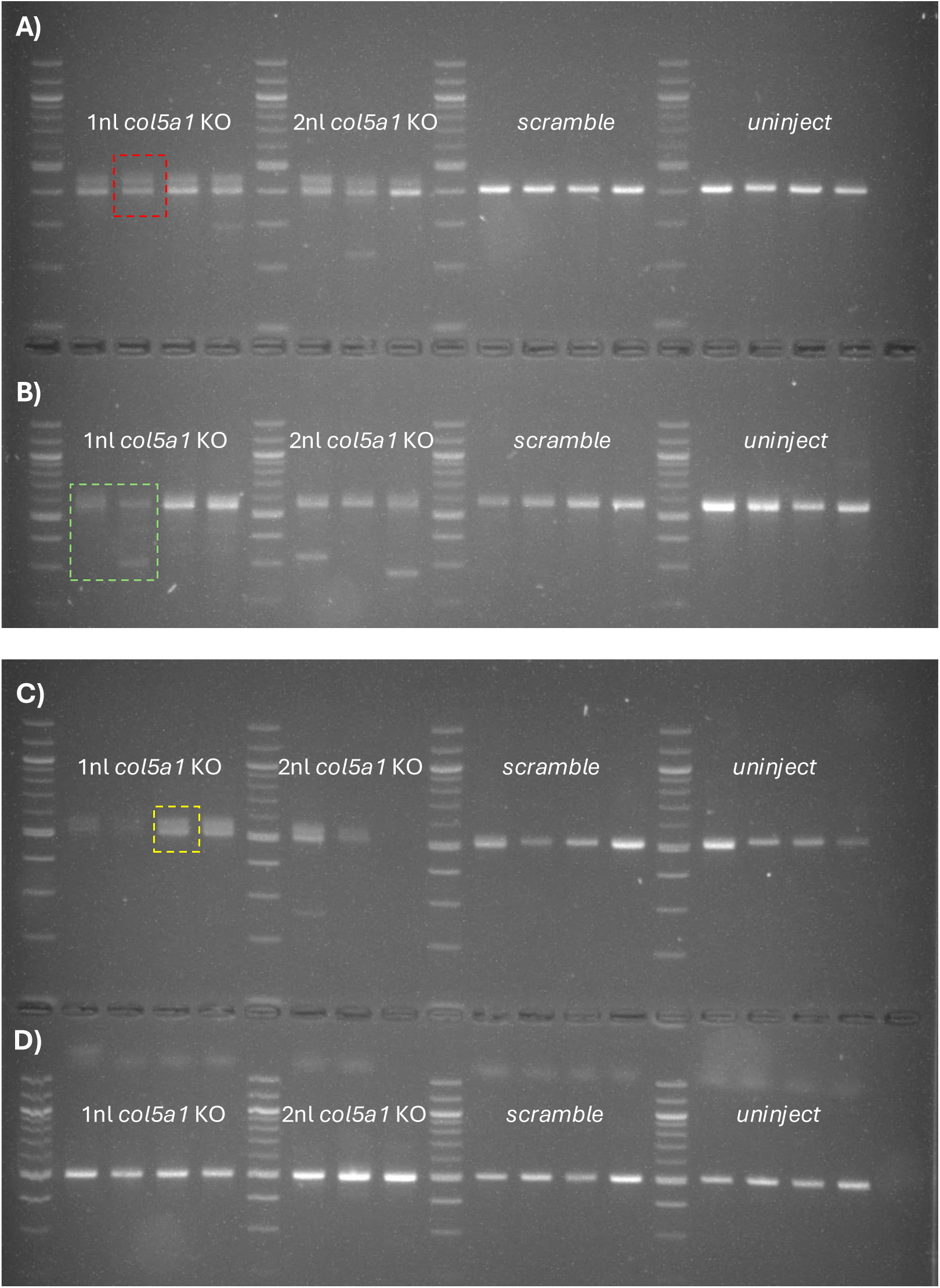
Representative gel electrophoresis of gene editing for *col5a1* exon targets by CRISPR-Cas9. Representative gel electrophoresis for *col5a1* exon targets with dosage tests for 1 and 2 nl of injected CRISPR-Cas9 gene editing reagents alongside uninject and *scramble* controls. **A.** Gel electrophoresis for exon 2 target, with red box highlighting multiple bands. **B.** Gel electrophoresis for exon 3 target, with green box highlighting multiple bands. **C.** Gel electrophoresis for exon 61 target, with yellow box highlighting multiple bands. **D.** Gel electrophoresis for exon 7 target.

**Suppl. Figure 3.**
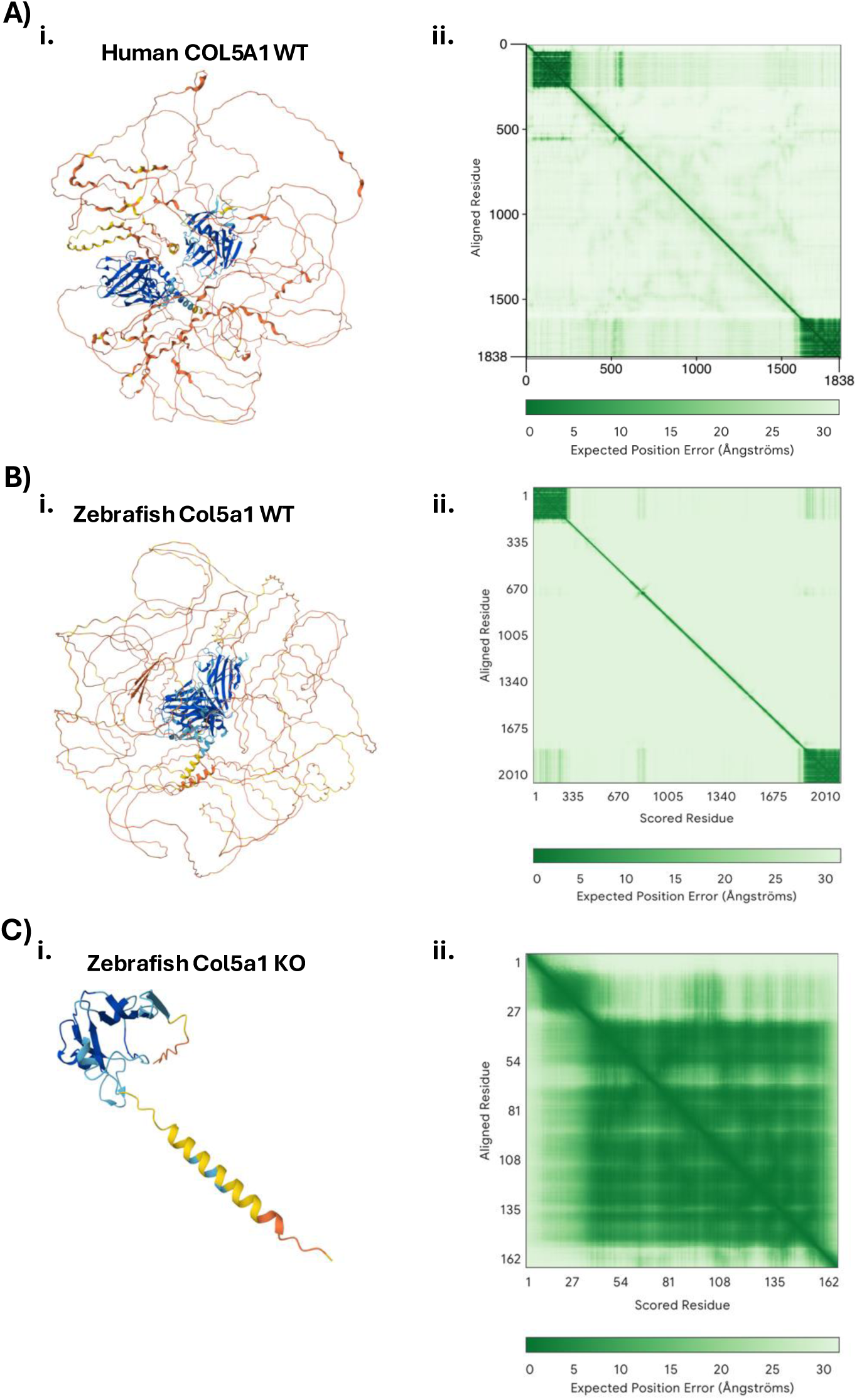
AlphaFold protein predictions for human and zebrafish type V collagen. **A.** AlphaFold prediction for wildtype human COL5A1 protein with **i)** 3D prediction image and **ii)** Predicted Aligned Error heatmap. **B.** AlphaFold prediction for wildtype zebrafish Col5a1 protein with **i)** 3D prediction image and **ii)** Predicted Aligned Error heatmap. **C.** AlphaFold prediction for knockout zebrafish Col5a1 protein generated from most frequently observed indels with **i)** 3D prediction image and **ii)** Predicted Aligned Error heatmap.

**Suppl. Figure 4.**
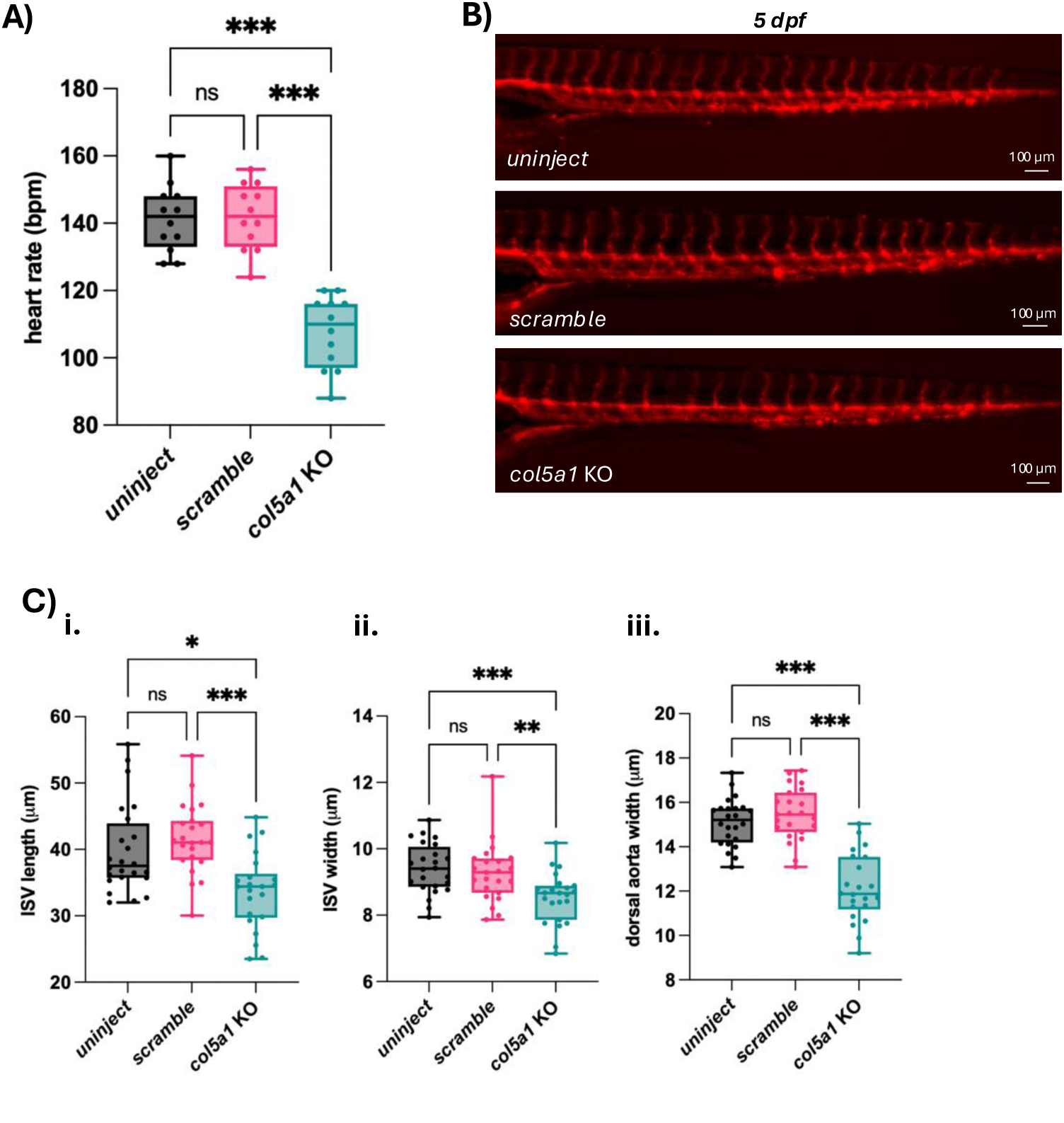
Cardiovascular phenotypes observed at 5 dpf. **A.** Heart rate quantification of experimental embryos at 5 dpf. Ordinary one-way ANOVA with Tukey’s multiple comparisons test performed. ns, not significant. *** *p* < 0.001. **B.** Representative immunofluorescent images of *Tg(kdrl:mCherry)* positive samples for uninject, *scramble*, and *col5a1* KO embryos at 5 dpf. Scale bars, 0.1 mm. **C.** Quantification of average **i)** ISV length, **ii)** ISV width, and **iii)** dorsal aorta width in each experimental condition at 2 dpf. Each dot in **(i)** represents the average of the total number of vessels found in the embryo tested. Each dot in **(ii)** and **(iii)** represents an average of 6 vessel measurements in an embryo. Kruskal-Wallis with Tukey’s multiple comparisons test performed for **(i)**. Lognormal ordinary one-way ANOVA with Tukey’s multiple comparisons test performed for **(ii)**. Ordinary one-way ANOVA with Tukey’s multiple comparisons test performed for **(iii)**. ns, not significant. *\* p* < 0.033. ** *p* < 0.002. *** *p* < 0.001.

## Notes

### Competing Interest Statement

The authors have declared no competing interest.

### Summary of Updates

Revised to ensure that bioRxiv headers do not overlap with the figure main body.

